# When Less Is Not More: DICEPro Mitigates the Impact of Incomplete Reference Matrices on Cellular Frequency Deconvolution

**DOI:** 10.64898/2026.06.17.732876

**Authors:** Kalidou Ba, Rodolphe Thiébaut, Xavier Hinaut, Boris Hejblum

## Abstract

Cellular deconvolution aims to estimate the frequencies of different cell populations from gene expression measurements in a biological sample. Supervised approaches, such as `CIBERSORTx` and `DISSECT`, critically depend on the reference signature matrix, which encodes the gene expression profiles of cell-types based on prior knowledge. Despite numerous deconvolution methods, the impact of missing cell populations in the reference matrix remains understudied. Here, we evaluate the robustness of state-of-the-art deconvolution approaches using simulations based on real dataset examples combined with statistical modeling, validated against published data, and multiple real benchmark datasets. Results show that deconvolution performance remains stable when the reference matrix includes most cell-types, but declines sharply as the matrix becomes incomplete, especially for abundant cell populations. To address the limitations of incomplete reference matrices, we introduce `DICEPro`, an optimization-based framework designed to enhance existing deconvolution methods. By systematically adjusting the reference signatures, `DICEPro` better accounts for missing or underrepresented cell populations, leading to improved precision and robustness. We show that `DICEPro` consistently boosts deconvolution performance across both simulated datasets, derived from real data examples, and multiple real biological datasets, offering a practical solution when standard methods are hindered by incomplete references.

## 1 Introduction

The cellular composition of biological tissues is diverse, as multiple cell-types coexist and interact within dynamic micro-environments. Understanding this heterogeneity is key to deciphering biological mechanisms. Cellular deconvolution has emerged as an essential computational tool enabling the estimation of cell-type frequencies from bulk RNA sequencing data [20, 25, 3]. Unlike single-cell RNA sequencing, which isolates individual cells, bulk RNA sequencing measures the aggregate expression of all cell-types present in a sample. Disentangling each cell-type contribution to gene expression through numerical deconvolution allows recovery of cell-type proportions without measuring them with specific assays such as flow cytometry.

Supervised deconvolution methods rely on predefined reference signature matrices encoding the gene expression profiles of known cell types. The precision of these methods critically depends on the quality, completeness, and biological relevance of these reference matrices [21, 15, 12]. In practice, biological variability, experimental conditions, and disease-specific contexts often create discrepancies between the cell populations represented in the reference and the actual sample composition. A particularly challenging scenario arises when specific cell populations are entirely absent from the reference matrix: such omissions can bias cell-type abundance estimates and degrade deconvolution performance [13]. This limitation is especially critical in specific tissues such as tumor microenvironments or in whole-blood disease contexts, where context-specific cell populations may be absent from standard reference matrices.

Despite the rapid growth of supervised deconvolution methods, the consequences of using incomplete reference matrices remain largely unexplored. The available approaches cover a broad methodological spectrum, from constrained regression and quadratic programming (with `ABIS` from Monaco et al. [18], `AutoGeneS` from Aliee and Theis [1], `DeconRNASeq` from Gong and Szustakowski [9], `EPIC` from Racle et al. [23], `LinDeconSeq` from Li et al. [16], or `QProg` and `QProgwc` from Gong et al. [10]) to robust and regularized regression (e.g. `CIBERSORTx` from Newman et al. [21], `DCQ` from Altboum et al. [2], `FARDEEP` from Hao et al. [11] or `RLR` and `RLS` from Sturm et al. [25]), to probabilistic or Bayesian modeling (e.g. `BayesPrism` from Chu et al. [6]), and to machine learning-based frameworks (such as `DISSECT` from Khatri et al. [14]).

Several benchmarking studies have systematically evaluated these methods under controlled conditions [25, 3, 19, 12], demonstrating that the algorithmic approach, the normalization strategy, and the selection of the reference matrix all influence independently the accuracy of the deconvolution. A large part of these evaluations have been conducted largely under the implicit assumption that the reference matrix adequately represents the cellular composition of the mixture. In practice, this assumption is frequently violated, particularly in histological or disease-specific contexts where certain cell populations may be absent or poorly represented in available references. Such mismatches can introduce substantial and systematic errors [13], but currently there is no established post-deconvolution correction strategy to address this limitation.

We introduce `DICEPro`, which stands for Deconvolution with Iterative Completion for Estimating cellular Proportion from RNA-seq data. This optimization-based framework for cellular deconvolution is designed to refine cell-type proportion estimates under incomplete reference matrices. Rather than functioning as a standalone deconvolution method, `DICEPro` operates as a post-deconvolution refinement layer that augments existing solutions by explicitly modeling the contribution of missing or underrepresented cell populations. The framework makes three original contributions. First, it extends the standard linear deconvolution model to account for populations absent from the reference, enabling proportion estimates to be corrected post hoc without modifying the upstream deconvolution pipeline. Second, it resolves the inherent tension between reconstruction accuracy and biological interpretability through a Pareto front strategy that makes the trade-off between these competing objectives explicit. Third, the framework is entirely agnostic to the choice of initialization method, accepting proportion estimates from any existing deconvolution tool as input and functioning as a general-purpose refinement layer broadly applicable across biological contexts and experimental platforms.

The remainder of this article is organized as follows. We first describe the `DICEPro` model and its optimization strategy, including the Pareto front approach used to balance residual fitting with biological constraints on mixing proportions in Methods. We continue to introduce realistic in silico simulations alongside our comparative assessment strategy to evaluate `DICEPro` and fourteen state-of-the-art deconvolution methods across scenarios of progressive reference incompleteness, as well as four benchmarking datasets derived from real bulk RNA-seq data with experimentally characterized cell-type proportions or in vitro biologically plausible cellular mixtures. Comparative performance is presented in Results, highlighting the consistent robustness gained from `DICEpro`. Finally, implications of our findings for practical deconvolution of bulk RNA-seq data are discussed in Discussion.

## 2 Methods

### 2.1 DICEPro

`DICEPro` takes as input an initial deconvoluted proportion matrix, obtained from any deconvolution algorithm, and improves it by explicitly modeling the additional contribution of missing cellular populations. The `DICEPro` framework is summarized in Fig. 1.

**Figure 1:**
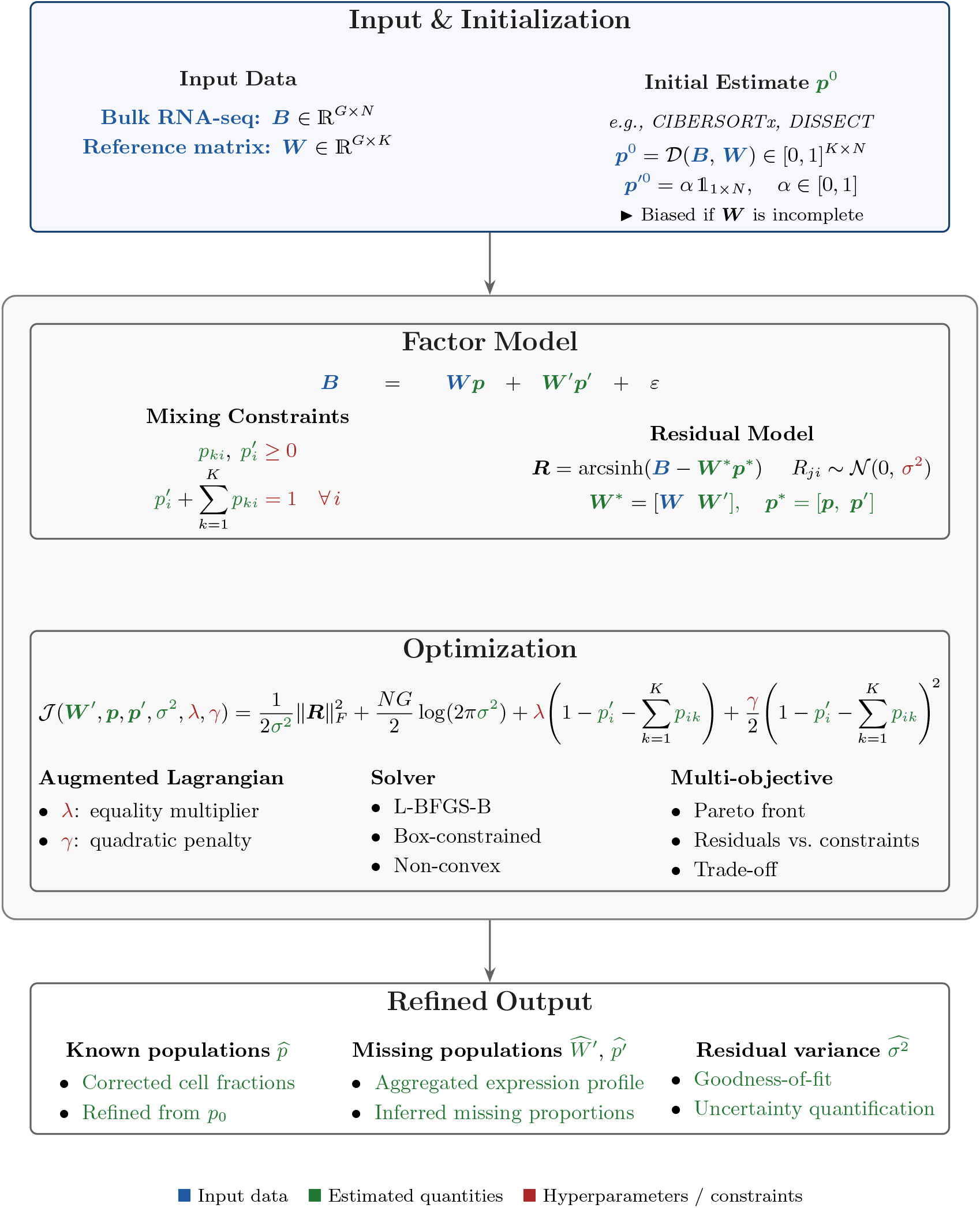
Overview of the DICEPro post-deconvolution refinement framework. `DICEPro` takes as input the proportion estimates produced by any deconvolution method and refines them to account for cell populations absent from the reference matrix, without requiring retraining or modification of the initial method.

#### 1. Factor model

Starting from bulk RNA-seq data ***B*** ∈ ℝ^*G×N*^ (where *G* is the number of genes and *N* the number of samples) and a (partial) reference matrix ***W*** ∈ ℝ^*G×K*^, an initial proportion matrix ***p***^0^ ∈ [0, 1]^*K×N*^ is first obtained from any reference-based deconvolution method. `DICEPro` then solves a constrained optimization problem that simultaneously refines the proportions ***p*** of known populations and infers the contribution of missing ones through two additional components: ***W*** ^*′*^ ∈ ℝ^*G×*1^, an aggregated expression profile of missing cell-types; and ***p***^*′*^ ∈ [0, 1]^1*×N*^, their corresponding proportions. The observed bulk expression matrix is now modeled as follows:

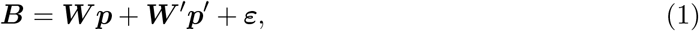

subject to constraints ensuring biological validity of the mixing proportions:

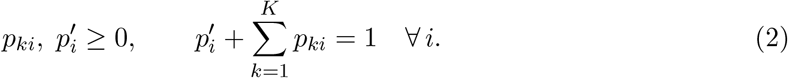

We further assume that the arcsinh-transformed residuals ***R*** = arcsinh(***B*** − ***W*** ^*\**^***p***^*\**^), with ***W*** ^*\**^ = [***W W*** ^*′*^] and ***p***^*\**^ = [***p p***^*′*^] combining known and missing components, each follow a Gaussian distribution *R*_*gi*_ ∼ *N* (0, *σ*^2^). This transformation stabilizes the variance and brings the residuals closer to this normality assumption [17], facilitating parameter estimation.

#### 2. Optimization

We minimize the corresponding negative Gaussian log-likelihood subject to the constraints in Equation (2) using an augmented Lagrangian formulation (See Supplementary Methods for the full derivation):

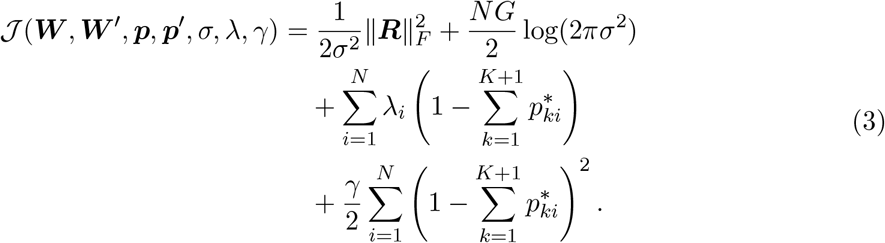

where *λ* is the Lagrange multiplier and *γ* is the quadratic penalty parameter. The resulting non-convex, box-constrained problem is solved with the L-BFGS-B algorithm [5, 27]. This optimization now involves two competing objectives: i) minimizing the Frobenius norm of the residuals and ii) satisfying the sum-to-one constraint. We therefore adopt a Pareto front strategy to navigate this trade-off: hyperparameters are explored via random search [4], and for each candidate combination, one Pareto-efficient solution is obtained. The selected solution achieves the best residual fit without violating the constraints (beyond an acceptable tolerance, see Supplementary Methods for details). This principled multi-objective approach [7] improves both the robustness and the interpretability of the deconvolution estimates.

#### 3. Initialization and outputs

`DICEPro` requires an initial proportion matrix to seed its constrained optimization. In this study, we used `CIBERSORTx` [21] for its stability and broad adoption across biological contexts. Crucially, however, the framework is agnostic to this choice: any valid proportion matrix can serve as an initial starting point, positioning `DICEPro` as a general-purpose post-deconvolution improvement layer rather than a method tied to a specific tool. The refined outputs of `DICEPro` are the corrected proportion matrix 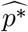, the aggregated missing-population signature 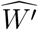, and the residual variance estimate 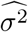 quantifying model un-certainty.

### Benchmarking datasets from real RNA-seq data

We benchmarked `DICEPro` against fourteen state-of-the-art supervised deconvolution methods on four datasets drawn from Nadel et al. [19], which together cover two mixture modalities and two tissue contexts. `CellMixtures` consists of 12 *in vitro* samples prepared by physical titration of six purified immune cell-types (B cells, CD4^+^ T cells, CD8^+^ T cells, monocytes, NK cells, and neutrophils) combined in known proportions. Expression was profiled on an Illumina HT12 BeadChip microarray, providing exact ground-truth fractions controlled at the time of sample preparation. `PBMC1NormMix` and `PBMC2NormMix` are two sets of 100 pseudo-bulk PBMC mixtures each, generated by aggregating single-cell RNA-seq profiles. PBMC1NormMix was derived from a previously published scRNA-seq dataset [19], while PBMC2NormMix was generated from publicly available PBMC data from 10x Genomics. Both cover seven immune cell-types: B cells, CD4^+^ T cells, CD8^+^ T cells, NK cells, monocytes, and neutrophils, with cell-type assignments validated against four to six marker genes per population (see Supplementary Figure S1 in Nadel et al. [19]). `StromalNormMix` consists of 100 pseudo-bulk stromal mixtures generated from 10x Genomics data and encompasses seven cell-types spanning immune and stromal compartments: B cells, CD4^+^ T cells, CD8^+^ T cells, macrophages, mast cells, endothelial cells, and fibroblasts. Cell-type assignment was originally performed by 10x Genomics. For all pseudo-bulk datasets, 1,000 cells were selected per sample, their expression values summed, and the resulting cell-type ratios recorded as ground-truth proportions. Together, these four datasets provide 312 samples with well-characterized cell-type composition, covering a broad range of immune and stromal cellular populations across two profiling platforms (microarray and pseudo-bulk from 10x scRNA-seq) and two proportion gold-standard strategies (experimental titration and *in silico* aggregation).

### Reference signature matrices

We evaluated eight reference matrices that cover diverse human tissues, cell-type granularities, and profiling platforms: `10XImmune`, `BlueCode`, `BluePrint-Blood`, `HPCA-Blood`, `HPCA-Stromal`, `ImmunoStates-Full`, `LM22`, and `SkinSignaturesV1`. The HPCA reference was subsetted into two tissue-appropriate versions: one restricted to seven blood cell-types (B cells, CD14^+^ monocytes, CD16^+^ monocytes, NK cells, neutrophils, CD4^+^ T cells, CD8^+^ T cells) and one covering six blood and stromal populations (B cells, CD4^+^ T cells, CD8^+^ T cells, endothelial cells, fibroblasts, macrophages). Some references include only a small set of filtered signature genes, while others span tens of thousands of genes captured by their respective technology. Prior to deconvolution, both bulk mixtures and reference profiles were jointly standardized by z-score normalization per gene across samples, following the preprocessing convention of CIBERSORT [20]. This diversity in gene number, cell-type compositions, as well as tissue contexts leads to varying deconvolution performance.

### Simulated datasets

Simulated bulk RNA-seq profiles were generated from the `BlueCode` reference matrix, which provides expression signatures for 34 cell subtypes organized into five major biological compartments: Immune, Stromal, Endothelial, Epithelial, and Muscle. This hierarchical organization enables a two-level Dirichlet sampling strategy that reflects the nested structure of cellular composition observed in complex tissues, producing pseudo-bulk samples with biologically coherent variability.

For each simulated sample, the group-level proportions ***π*** = (*π*_1_, …, *π*_5_) were first drawn from a Dirichlet distribution, ***π*** ∼ Dir(*α*_comp_), where *α*_comp_ was calibrated to match the variability between-samples observed in experimental bulk RNA-sequencing datasets [19]. Within each compartment *c*, subtype-level proportions 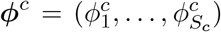 were then sampled from a second Dirichlet distribution, ***ϕ***^*c*^ ∼ Dir 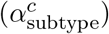, where *S*_*c*_ denotes the number of subtypes in the compartment *c* and 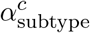 encodes heterogeneity within each group. The final proportion vector was obtained by compositional rescaling 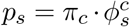, ensuring that all 36 subtypes satisfy simplex constraints.

The bulk expression profile ***b*** ∈ ℝ^*G*^ for each simulated sample was calculated as a linear mixture of reference signatures: ***b*** = ***Wp***. All expression values were retained on the linear scale prior to noise addition, consistent with the modeling assumptions of `DICEPro` and other standard reference-based deconvolution frameworks [3].

To emulate the variability observed in real bulk RNA-seq data, two independent noise components were introduced. First, biological variability was modeled as multiplicative perturbations of the proportions of compositional cells, where each component was transformed as

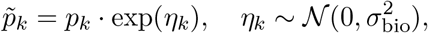

followed by renormalization to enforce simplex constraints. The parameter σ_bio_ was fixed to induce moderate variability between the samples consistent with the experimentally observed heterogeneity. Second, technical variability was modeled as additive noise in the gene proportional to the magnitude of the expression, where observed expression values were generated as follows:

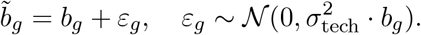

The parameter 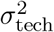 was chosen to match the coefficient of variation observed in real datasets, and negative values resulting from additive noise were truncated to zero to preserve biological interpretability.

### 2.2 Comparative assessment strategy

All methods were evaluated on the real and simulated datasets described above, using experimentally determined or construction-derived cell-type fractions as ground truth. To assess robustness to reference incompleteness, we leveraged simulated data by systematically removing predefined groups of cell types (first B cells, then CD4^+^ T cells, CD8^+^ T cells, NK cells, myeloid cells, and finally endothelial cells) either individually or cumulatively. In each scenario, deconvolution was performed using the reduced reference matrix, and predictions were compared to true proportions across 50 Monte Carlo replicates.

Performance was assessed using two complementary metrics. The Pearson correlation coefficient (*r*) was computed sample-by-sample and averaged across samples, providing a widely used measure of concordance in previous deconvolution benchmarks [25, 3]. However, because proportion estimates exhibit a hierarchical structure across samples and cell types, treating observations as independent may lead to biased performance estimates [8]. To address this, we additionally report the hierarchical relative Root Mean Square Error (*hrRMSE*), derived from a linear mixed-effects model with nested random effects that partition variance into biological signal and estimation error. Unlike the standard *RMSE, hrRMSE* normalizes the estimation error by the total biological variance, allowing fair comparison across datasets and cell types with heterogeneous abundance levels. Full model specification is provided in the Supplementary.

## 3. Results

### 3.1 Performance under complete reference conditions

We first evaluated deconvolution performance under idealized conditions where the reference matrix fully represented all cell types present in the mixture. Under these settings, the benchmarked methods showed heterogeneous accuracy (Fig. 2A), separating into three categories of performance. Robust regression-based approaches achieved the highest baseline performance, with `FARDEEP` leading across both metrics, while `BayesPrism` performed comparably while relying on a probabilistic framework. A second tier comprising `LinDeconSeq`, `RLR`, `RLS`, `CIBERSORTx`, and `ABIS` achieved intermediate accuracy. In contrast, marker-based methods such as `EPIC` performed substantially worse, consistent with their known sensitivity to normalization [3]. Initialized from `CIBERSORTx` estimates, `DICEPro` reached accuracy levels comparable to the top-performing methods, and systematically improved upon its initializer in both *r* and *hrRMSE* in each Monte Carlo replicate. This indicates that the post-deconvolution optimization step effectively reduces residual noise and estimation bias, even when the reference matrix is complete (Fig. 2C–D).

**Figure 2:**
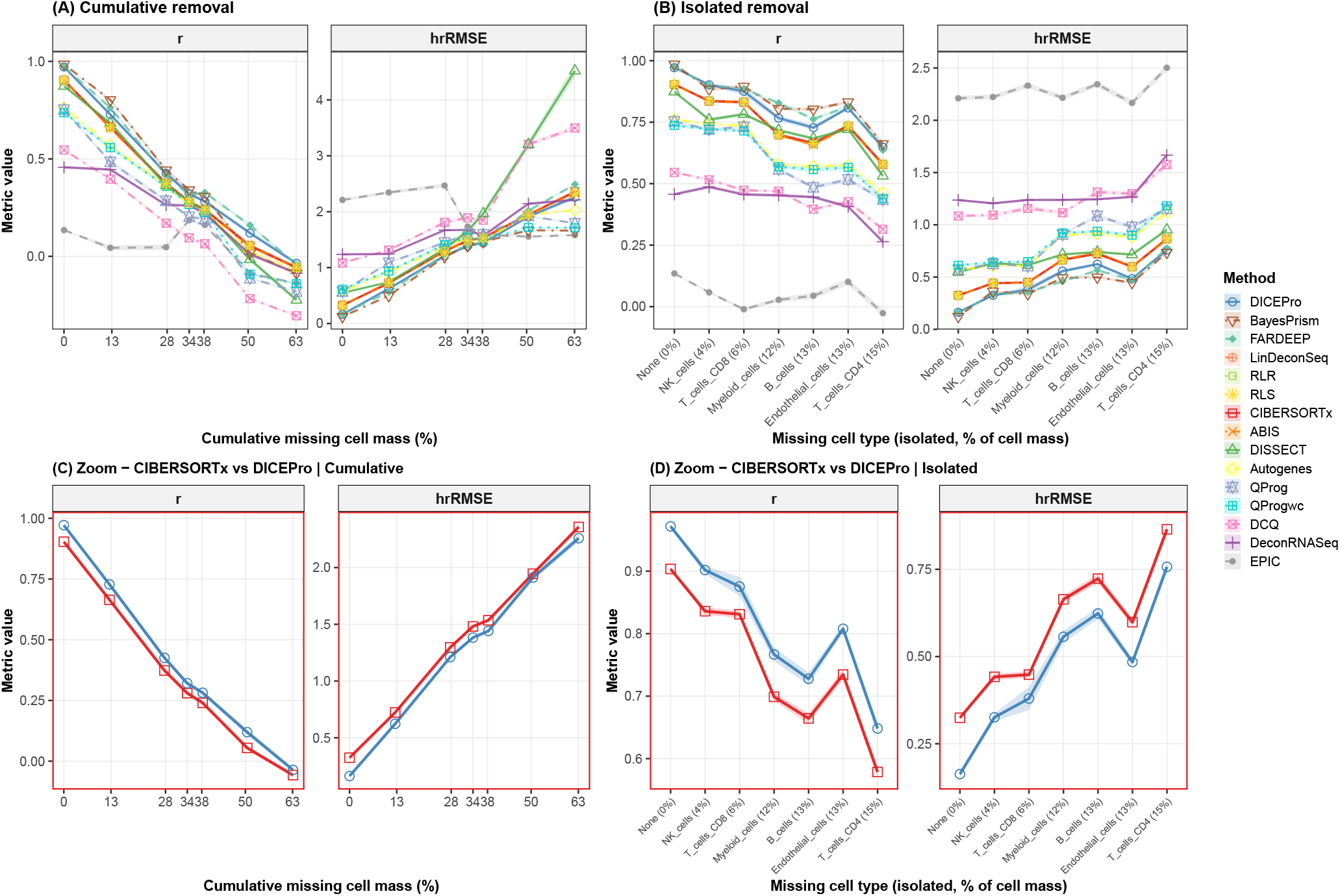
Performance of DICEPro and fourteen comparator cellular deconvolution methods under cumulative and isolated missing cell-type scenarios in simulated bulk RNA-seq data. Panels (A) and (B) display mean ± SE Pearson correlation (*r*) and Hierarchical Relative Root Mean Square Error (*hrRMSE*) across 50 Monte Carlo replicates under cumulative (A) and isolated (B) cell-type removal scenarios. In (A), cell types were removed in order of increasing abundance, corresponding to a cumulative missing cell mass from 0% to ∼ 63%. In (B), each cell type (B cells, CD4^+^ T cells, CD8^+^ T cells, NK cells, myeloid cells, and endothelial cells) was removed individually, with the percentage of removed cell mass indicated on the x-axis. Panels (C) and (D) show the same scenarios restricted to `CIBERSORTx` and `DICEPro` (red-bordered panels), highlighting the gain attributable to post-deconvolution optimization.

### 3.2 Robustness to incomplete reference matrices

We assessed robustness to ***W*** incompleteness by progressively removing cell types, thus simulating increasing levels of missing cell mass.

### 3.2.1 Cumulative removal

Cell types were removed sequentially by group (B, T_CD4, T_CD8, NK, Myeloid, Endothelial, Fibroblast, Smooth_muscle), generating scenarios ranging from complete reference coverage to approximately 63% missing cell mass (Fig. 2A). All methods exhibited a monotonic decrease in performance as incompleteness increased, but the rate of degradation differed markedly between algorithmic families. Methods that performed best under complete conditions, including BayesPrism and FARDEEP, showed the steepest decline, with their advantage diminishing at intermediate levels of missingness. At approximately 38% of missing mass, `DICEPro` stopped showing a significant advantage in performance over these methods. Beyond roughly 50% of missing mass, all methods had near-zero or negative correlations, reflecting a fundamental identifiability limit of linear deconvolution rather than a limitation specific to any individual approach. Across all levels of incompleteness, `DICEPro` consistently outperformed its `CIBERSORTx` initializer. This suggests that its correctly specified mixture model and the Pareto-front optimization framework successfully increase robustness to reference degradation.

#### 3.2.2 Isolated removal

To disentangle the effect of missing cell identity from that of cumulative missingness, we next removed individual cell-types in isolation (Fig. 2B). Performance loss depended not only on the abundance of the missing population but also on its transcriptional distinctiveness and co-expression with remaining cell-types. The removal of transcriptionally distinct populations, such as NK cells, despite their relatively low abundance, resulted in disproportionately higher degradation of performance. In contrast, the removal of populations with more common expression profiles produced more moderate effects, even at higher abundance levels. Across all scenarios, `DICEPro` consistently ranked among the top-performing methods and improved upon its `CIBERSORTx` initialization (Fig. 2C and 2D). Even with differnt algorithm initializations, `DICEPro` performance remained very stable and comparable to the top performing approaches (See Supplementary Fig. S1 and S2).

### 3.3 Validation on real bulk RNA-seq datasets

To validate these findings under experimentally grounded conditions, all methods were evaluated on four independent bulk RNA-seq datasets using eight reference matrices of varying composition and cell-type coverage (Fig. 3). Across the resulting 32 dataset-reference combinations, `DICEPro` achieved the highest mean performance and outperformed `CIBERSORTx` in the vast majority of scenarios, ranking first in nearly half of all combinations (17 out of 32, i.e., 53%) and improving upon its initializer in most (72%) cases. Compared with the broader benchmark landscape, `DICEPro` also outperformed `LinDeconSeq`, `FARDEEP`, and `BayesPrism` on average, with the most consistent advantage observed under conditions of reference-sample mismatch.

**Figure 3:**
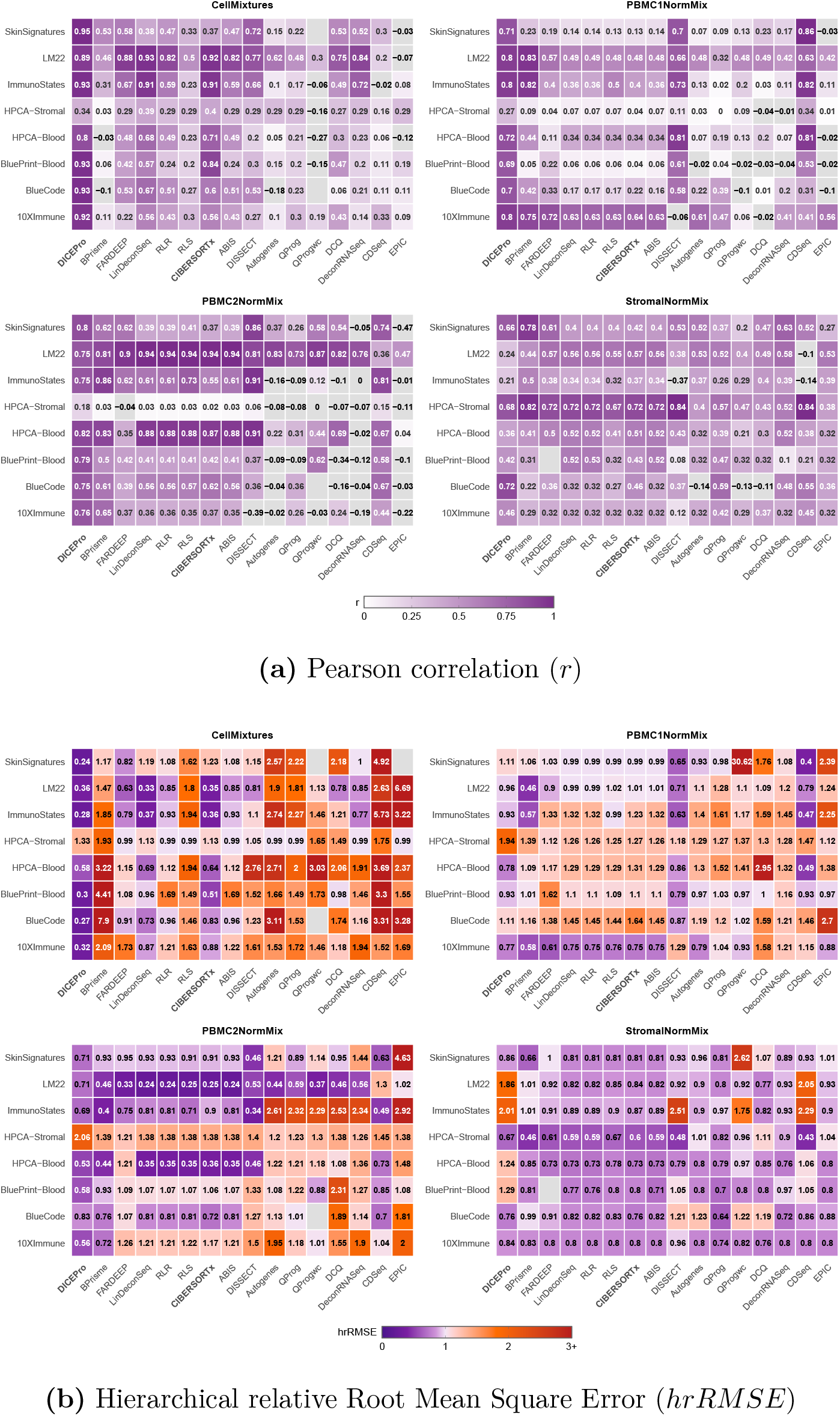
Deconvolution performance on real bulk RNA-seq datasets. Heatmaps of (a) Pearson correlation (*r*) and (b) hierarchical Relative Root Mean Square Error (*hrRMSE*) for all benchmark methods across four datasets (*CellMixtures, PBMC1NormMix, PBMC2NormMix*, and *StromalNormMix*) and eight reference matrices. In panel (a), the color scale ranges from white (low positive *r*, poorer performance) to dark purple (high positive *r*, better performance). Grey cells indicate negative correlations (*r <* 0) or missing/undefined values. In panel (b), values below 1 transition from dark violet (closer to 0, better performance) through light purple, whereas values above 1 transition from orange to dark red (poorer performance). For visualization purposes, values greater than 3 are truncated to 3. Grey cells in panel (b) indicate missing or undefined values.

Performance gains were particularly pronounced in the `CellMixtures` and `PBMC` datasets, where reference incompatibility strongly penalized competitive methods. In these settings, `DICEPro` demonstrated substantial robustness, with the largest improvements observed when tissue-inappropriate references were used — precisely the scenarios in which missing cell populations are most likely to occur.

The `StromalNormMix` dataset represented a more challenging and structured context. With tissue-matched references, methods explicitly modeling biological structure, such as `BayesPrism`, achieved the highest performance, reflecting the advantage of probabilistic approaches under well-specified conditions. In this setting, `DICEPro` showed slightly lower performance for specific reference configurations but remained competitive and recovered strong accuracy when alternative references were used. This variability once again highlights the crucial role of reference quality in deconvolution performance.

### 3.4 Overall assessment

Together, these results establish two consistent findings across both simulated and real datasets. First, reference incompleteness systematically degrades deconvolution accuracy across all methods, with the magnitude of degradation jointly depending on the fraction of missing cell mass and on the distinctiveness of the expression profile the absent populations. Second, `DICEPro` consistently improves upon its `CIBERSORTx` initialization and maintains competitive or superior performance across a wide range of conditions. Notably, the magnitude of improvement increases with the degree of reference mismatch, highlighting the specific advantage of post-deconvolution refinement in less-than-ideal reference signature settings.

Overall, these findings position `DICEPro` as a robust and effective refinement strategy for deconvolution analyzes in realistic scenarios where reference matrices are inherently incomplete or imperfect. The biological implications of these results are discussed in Discussion.

## 4 Discussion

`DICEPro` demonstrated consistent improvements over its `CIBERSORTx` initialization across both simulated and real bulk RNA-seq datasets, establishing post-deconvolution optimization as a practically viable strategy to refine cell-type proportion estimates. Its open-source implementation ensures full reproducibility of the analyzes presented here and provides a transparent foundation for future methodological developments, providing a transparent foundation for future methodological developments (in contrast to widely adopted platforms such as `CIBERSORTx`, whose partially closed-source implementation limits independent reproduction and community-driven extension).

Across all benchmark conditions, the quality and completeness of the reference signature matrix emerged as the dominant factor governing deconvolution accuracy, overshadowing algorithmic differences between methods. In real datasets, performance varied substantially across the eight reference matrices tested, with tissue-matched references producing markedly higher *r* than mismatched alternatives. This confirms that no optimization strategy, however sophisticated, can fully compensate for a fundamentally misspecified reference. Practically, this underscores reference selection as a critical upstream decision in any deconvolution workflow: investing in reference curation is likely to yield larger precision gains than switching between deconvolution algorithms. `DICEPro` does not eliminate this constraint, but provides a principled mechanism to partially mitigate its effects by reallocating a fraction of the expression away from available reference types when some populations are absent.

Although simulation results are, by construction, free from technical variability, performance on real bulk RNA-seq datasets was substantially influenced by normalization choices. `DICEPro`’s objective function is derived from a Gaussian likelihood framework, which explicitly assumes that residuals are normally distributed with constant variance. This assumption directly conditions the validity of the proportion estimates: the optimization is only well-founded when this normality condition holds. Normalization of both the mixture samples and the reference matrix is therefore an important requirement of the model rather than an optional preprocessing step, and normalization incompatibility between reference and mixture can be an important driver of performance variability on real datasets.

A recurring challenge in deconvolution benchmarking is the choice of evaluation metric. Standard metrics such as Pearson correlation or *RMSE* treat all proportion estimates as independent observations, ignoring the nested structure of deconvolution data (where measurements are jointly organized by samples and cell populations) which can produce biased or over-optimistic assessments. The hierarchical mixed-effects framework used to derive *hrRMSE* in this study provides a more principled foundation, explicitly partitioning variance into biological signal and estimation error. This framework naturally extends to complementary indicators such as the intraclass correlation coefficient (*ICC*) and the concordance correlation coefficient (*CCC*), which can be derived from the same variance decomposition and provide complementary perspectives on deconvolution performance (see Supplementary Section S3). Metric choice should therefore be treated as a first-class consideration in benchmarking studies, alongside algorithmic and reference design decisions.

There are three main limitations to the current framework. First, although `DICEPro` demonstrated resilience to partial reference incompleteness, performance degraded substantially when cumulative missing cell mass exceeded approximately 50%. Beyond this threshold, the optimization becomes ill-conditioned: the remaining reference types cannot adequately span the mixture space, and the constrained optimization redistributes the signal into increasingly unreliable estimates. This reflects a fundamental identifiability limitation shared across all de-convolution methods rather than a limitation specific to `DICEPro`. Second, `DICEPro` inherits the structural properties of its initialization. When the initializer produces severely biased estimates, as observed with strongly mismatched references on the `StromalNormMix` dataset, the optimization may converge to a local minimum close to the initial solution, limiting the improvement. Selecting a better initializer or incorporating a multi-start strategy drawing from several independent initializers could substantially alleviate this dependency and extend the practical applicability of the framework. Third, reconstruction error alone — as measured by the Frobenius norm — is insufficient to fully characterize the optimization objective. Minimizing reconstruction error without constraints can drive proportion estimates outside the biological simplex, yielding numerically accurate but biologically implausible solutions. `DICEPro` addresses this issue by treating reconstruction fidelity and the unit-sum constraint as separate objectives, and explicitly balancing them through a Pareto front construction (See Supplementary Fig. S3). The resulting front retains non-dominated solutions, from which a knee-point criterion provides a principled default operating point. Importantly, this formulation makes the trade-off explicit rather than relying on a single scalar penalty parameter, allowing different operating points to be selected depending on the biological context.

`DICEPro` enforces the simplex constraint through an augmented Lagrangian formulation that jointly minimizes reconstruction error while penalizing unit-sum violations, providing a rigorous link between the biological requirement for proportion interpretability and the structure of the objective function. The inner subproblem is currently solved with L-BFGS-B, a quasi-Newton method that exploits gradient information and efficiently handles bound constraints. Alternative optimization strategies may further improve convergence and scalability.

On the evaluation side, the Pareto front construction currently treats the reference signature matrix as fixed, whereas reference signatures are in practice subject to biological and technical variability. Incorporating uncertainty quantification, for instance through sampling of reference signatures, would allow `DICEPro` to propagate this uncertainty into the final estimates, providing confidence intervals alongside point estimates and further strengthening the framework across diverse reference construction strategies.

The results of this study support several practical recommendations for deconvolution workflows. In particular, reference signature selection and normalization compatibility should be carefully evaluated prior to method choice, as they can have a major impact on performance, often comparable to or exceeding differences between algorithms. When the reference matrix is incomplete or partially mismatched, post-deconvolution refinement with `DICEPro` provides a principled and computationally efficient correction layer that can be applied to any initializing method.

Beyond performance, reproducibility and methodological transparency are essential requirements for computational tools intended for broad scientific use [24, 26]. While `CIBERSORTx` is widely used and effective, its partially closed-source implementation can limit its deployment, independent reproducibility and methodological extensions [21]. In contrast, `DICEPro` will be released as an open-source R package, fully documented and publicly available on GitHub (https://github.com/kalidouBA/dicepro), enabling transparent and reproducible analyzes. Given the sensitivity of deconvolution results to implementation details and preprocessing choices [3], such transparency is critical for reproducibility and reliable benchmarking [22].

## Supporting information

Supplementary materials for "When Less Is Not More: DICEPro Mitigates the Impact of Incomplete Reference Matrices on Cellular Frequency Deconvolution"

## 7 Acknowledgments

The authors thank Bastien Chassagnol for interesting discussions about cellular deconvolution and numerical optimisation.

