## Supplementary materials for "When Less Is Not More: DICEPro Mitigates the Impact of Incomplete Reference Matrices on Cellular Frequency Deconvolution" for "When Less Is Not More: DICEPro Mitigates the Impact of Incomplete Reference Matrices on Cellular Frequency Deconvolution"

**S1 DICEPro methodological framework** **1**

**S3 Hierarchical performance metrics for deconvolution evaluation** **14**

### S1 DICEPro methodological framework

Theoretical method for optimizing the parameters matrix decomposition under the given constraints and regularization terms using gradient descent.

$$B = Wp + W'p' + \varepsilon$$

$$\text{s.t.} \quad \begin{cases} p_{i,j},\, p'_{i,j} \geq 0 \\ \|p_i + p'_i\|_1 = 1 \end{cases}$$

- $B \in \mathbb{R}^{G\times N}$ is the observed gene expression matrix.
- $W \in \mathbb{R}^{G\times K}$ and $W' \in \mathbb{R}^{G\times K'}$ are reference expression matrices.
- $p \in \mathbb{R}^{K\times N}$ and $p' \in \mathbb{R}^{K'\times N}$ are the mixing proportion matrices.
- $\varepsilon$ represents the residual error matrix.
- We consider $K' = 1$, so that $W'$ and $p'$ reduce to vectors.

#### S1.1 Derivation of the Objective Function

##### Gaussian Assumption

The residuals are defined as:

$$R = \text{arcsinh}(B - W^* p^*)$$

where:

- $W^* = [W \mid W'] \in \mathbb{R}^{G\times K^*}$ is the combined reference matrix.
- $p^* = [p^\top \mid p'^\top]^\top \in \mathbb{R}^{K^*\times N}$ is the combined proportion matrix.
- $K^* = K + K'$.

We assume that the residuals follow a Gaussian distribution with mean 0 and variance $\sigma^2$:

$$R_{ij} \overset{\text{i.i.d.}}{\sim} \mathcal{N}(0, \sigma^2)$$

##### Likelihood Function

The likelihood of observing $B$ given $W^*$, $p^*$, and $\sigma^2$ is:

$$\mathcal{L}(B \mid W^*, p^*, \sigma^2) = \prod_{i,j} \frac{1}{\sqrt{2\pi\sigma^2}} \exp\left(-\frac{R_{ij}^2}{2\sigma^2}\right)$$

Taking the natural logarithm yields the log-likelihood:

$$\log \mathcal{L}(B \mid W^*, p^*, \sigma^2) = -\frac{\|R\|_F^2}{2\sigma^2} - \frac{NG}{2}\log(2\pi\sigma^2)$$

where $N$ is the number of samples and $G$ is the number of genes.

##### Augmented Lagrange multiplier

To enforce the simplex constraint $\sum_{k=1}^{K^*} p_{ki}^* = 1$ for each sample $i$, we leverage an augmented Lagrange multiplier yielding the two additional terms below:

$$\lambda \sum_{i=1}^{N} \left| 1 - \sum_{k=1}^{K^*} p_{ki}^* \right| \quad \text{and} \quad \frac{\gamma}{2} \sum_{i=1}^{N} \left( 1 - \sum_{k=1}^{K^*} p_{ki}^* \right)^2$$

#### S1.2 Objective Function

Combining the negative log-likelihood with the regularization terms, the objective function to minimize is:

$$\mathcal{J}(W^*, p^*, \sigma, \lambda, \gamma) = \frac{\|R\|_F^2}{2\sigma^2} + \frac{NG}{2}\log(2\pi\sigma^2) + \lambda \sum_{i=1}^{N} \left| 1 - \sum_{k=1}^{K^*} p^*_{ki} \right| + \frac{\gamma}{2} \sum_{i=1}^{N} \left( 1 - \sum_{k=1}^{K^*} p^*_{ki} \right)^2$$

##### Initialization

- Initialize $W^*$ as $[W \mid W']$, where the $K$ columns of $W \in \mathbb{R}^{G \times K}$ are held **fixed** throughout optimization, and $W' \in \mathbb{R}^{G \times 1}$ is initialized as a vector of ones representing the unknown cell-type component.
- Initialize $p^*$ as $[p^\top \mid p'^\top]^\top$, where $p \in \mathbb{R}^{K \times N}$ contains the cell-type proportions estimated by `CIBERSORTx` on the incomplete reference $W$, and $p' \in \mathbb{R}^{1 \times N}$ is a row vector whose constant value is sampled uniformly from $[0, 1]$ via random search and replicated across all samples.
- Initialize $\lambda$ and $\gamma$ via random search, each sampled log-uniformly on $[1, 10^5]$.
- Initialize $\sigma$ as the mean standard deviation of the residuals $R$.

##### Gradient Derivations

Let $\delta = \operatorname{arcsinh}(B - W^*p^*)$ and $\delta' = \frac{1}{\sqrt{1+(B-W^*p^*)^2}}$ denote the elementwise derivative of arcsinh evaluated at $B - W^*p^*$. By the chain rule:

- **Gradient with respect to $W'$:**

$$\frac{\partial \mathcal{J}}{\partial W'} = -\frac{1}{\sigma^2} (R \odot \delta') \, p'^\top$$

  The first $K$ columns of $W^*$ (corresponding to $W$) are kept fixed and are not updated.

- **Gradient with respect to $p^*$:**

$$\frac{\partial \mathcal{J}}{\partial p^*} = -\frac{1}{\sigma^2} W^{*\top}(R \odot \delta') - \lambda \operatorname{sign}\left( 1 - \sum_{k=1}^{K^*} p^*_{ki} \right) \mathbf{1}^\top - \gamma \sum_{i=1}^{N} \left( 1 - \sum_{k=1}^{K^*} p^*_{ki} \right) \mathbf{1}^\top$$

  where $\odot$ denotes elementwise multiplication. A subgradient is used for the $\ell_1$ term when $1 - \sum_k p^*_{ki} = 0$.

- **Gradient with respect to $\sigma$:**

$$\frac{\partial \mathcal{J}}{\partial \sigma} = -\frac{\|R\|_F^2}{\sigma^3} + \frac{NG}{\sigma}$$

- **Gradient with respect to $\lambda$:**

$$\frac{\partial \mathcal{J}}{\partial \lambda} = \sum_{i=1}^{N} \left| 1 - \sum_{k=1}^{K^*} p^*_{ki} \right|$$

- **Gradient with respect to $\gamma$:**

$$\frac{\partial \mathcal{J}}{\partial \gamma} = \frac{1}{2} \sum_{i=1}^{N} \left( 1 - \sum_{k=1}^{K^*} p^*_{ki} \right)^2$$

#### S1.3 Empirical Validation of the Solver and Penalty-Based Formulation

All empirical results presented in this section (namely the solver benchmark presented in Figure S1) and the constraint enforcement analysis presented in Figure S2) are demonstrated using real-world dataset: bulk RNA-seq data from the **CellMixtures** dataset (Nadel et al., 2021), a collection of in vitro cellular mixtures with experimentally characterised cell-type proportions that serve as ground truth. The reference signature matrix used was **LM22** (Newman et al., 2015), a widely adopted 22-cell-type immune signature matrix. Gene expression values were z-score normalised per gene prior to deconvolution, and initial proportion estimates were obtained from `CIBERSORTx` (Newman et al., 2019) run on the LM22 reference.

##### S1.3.1 Choice of the L-BFGS-B Optimization Algorithm

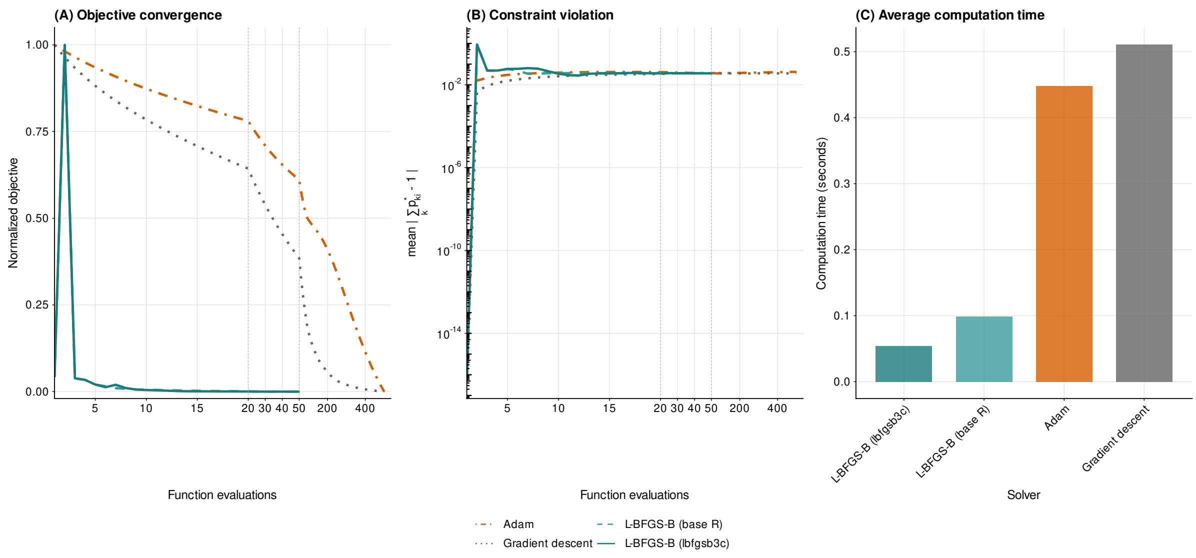

Figure S1: **Empirical comparison of four optimisation solvers applied to the `DICEPro` objective function on the CellMixtures and LM22 benchmark dataset.** Panel (A) shows the normalised objective value $\mathcal{J}$ as a function of the number of function evaluations for each solver. The x-axis uses a piecewise-linear scale with a hinge at iteration 50 to preserve early-convergence detail while displaying the full trajectory up to the maximum iteration budget. Panel (B) shows the mean constraint violation $\overline{|c_i|} = N^{-1}\sum_{i=1}^{N}|1-\sum_k p^*_{ki}|$ on a logarithmic scale. The near-zero violation at the first evaluation reflects the normalised initialisation of $\boldsymbol{p}^*$; the subsequent spike corresponds to the first gradient step away from this initial point. Panel (C) compares total computation time across solvers under identical initialisation conditions.

We compared L-BFGS-B (`lbfgsb3c` and base-`R` implementations) against two representative first-order methods Adam (Kingma and Ba, 2015) and projected gradient descent with Armijo backtracking on the bulk CellMixtures and LM22 benchmark dataset. Results are summarised in Figure S1.

Panel (A) shows that both L-BFGS-B implementations reach a stable minimum in fewer than 10 iterations, using substantially fewer function evaluations than first-order methods, reflecting the curvature information captured by the limited-memory Hessian approximation. Panel (B) shows that L-BFGS-B reaches a feasible region within the first few evaluations and maintains a stable constraint violation thereafter, while gradient descent requires the full iteration budget to approach the same level. Adam exhibits a persistently higher constraint violation, reflecting a trade-off between likelihood minimisation and simplex feasibility: although Adam converges to a lower objective value, this comes at the cost of reduced constraint satisfaction. Panel (C) compares the average computation time across optimizers. The `L-BFGS-B (lbfgsb3c)` implementation achieves the lowest runtime (0.054 s on average), followed by the base R implementation of `L-BFGS-B` (0.099 s). First-order methods such as Adam (0.448 s) and gradient descent (0.511 s) are substantially slower due to the large number of iterations required to reach convergence (500 iterations). Despite these differences in runtime, all solvers reach comparable feasibility levels, with final constraint violations on the order of $10^{-2}$ to $10^{-1}$.

Beyond these empirical observations, L-BFGS-B provides two additional structural advantages for the `DICEPro` problem. First, the non-negativity constraints $p^*_{ki} \geq 0$ are handled natively via gradient projection onto the active bounds (Byrd et al., 1995), eliminating external projection steps. Second, the limited-memory Hessian approximation reduces memory and computational complexity from $\mathcal{O}(n^2)$ for exact Newton methods to $\mathcal{O}(mn)$ with $m \ll n$ (Nocedal, 1980; Liu and Nocedal, 1989), making the approach scalable to genome-wide applications.

##### S1.3.2 Numerical Validation of the Augmented Lagrangian Formulation

**Gradient correctness (panel A).** Analytical gradients with respect to $\boldsymbol{W}'$ and $\boldsymbol{p}^*$ were verified against central finite differences ($\varepsilon = 10^{-5}$) on 30 randomly sampled coordinates of $\boldsymbol{\theta}$. As shown in panel A, gradients for $\partial\mathcal{J}/\partial\boldsymbol{W}'$ remain moderate in magnitude (e.g., $-4.38 \times 10^{-1}$ to $+3.89 \times 10^{-1}$) while those for $\partial\mathcal{J}/\partial\boldsymbol{p}^*$ exhibit a distinctly wider dynamic range (e.g., $-1.54 \times 10^2$ to $+2.00 \times 10^2$). Across all sampled coordinates, relative errors between analytical and numerical gradients are consistently below $6 \times 10^{-6}$, with most errors on the order of $10^{-7}$ to $10^{-9}$. This confirms the correctness of the analytical gradient expressions for both parameter types.

**Penalty landscapes (panel B).** The plots show the regime $c_i \geq 0$, corresponding to positive constraint violations. In this regime, the linear term increases proportionally with the violation magnitude, while the quadratic term induces a smooth convex growth. For representative parameter values, the linear term dominates under large violations, whereas the quadratic term provides stronger curvature near feasibility. This highlights the complementary roles of both components: robust constraint enforcement versus smooth convergence in the vicinity of the feasible region.

#### S1.4 Robustness of `DICEPro` to the choice of deconvolution method for initialization

Supplementary Figures S3 and S4 summarize the performance of the method under cumulative and isolated missing cell type conditions, evaluated using the Pearson correlation coefficient ($r$) and the hierarchical relative root mean square error ($hrRMSE$). All methods exhibited a progressive decline in performance as the proportion of missing cell mass increased, with the magnitude of degradation reflecting the cumulative loss of reference information available to the deconvolution algorithm. This degradation was observed

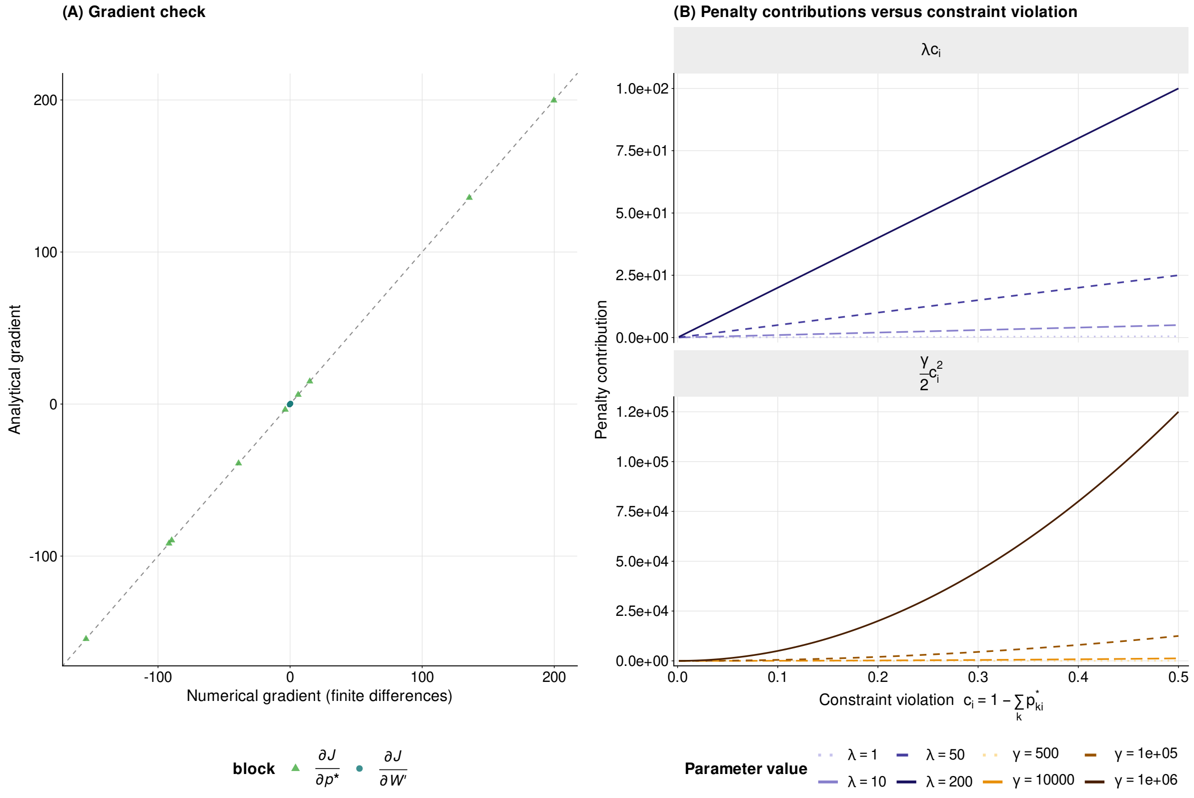

Figure S2: **Numerical validation of the augmented Lagrangian formulation underlying `DICEPro`, applied to the CellMixtures–LM22 benchmark dataset.** Panel (A) compares analytical gradients against numerical gradients computed by central finite differences ($\varepsilon = 10^{-5}$) on 30 randomly sampled coordinates of $\boldsymbol{\theta}$, stratified by parameter block ($\partial \mathcal{J}/\partial \boldsymbol{W}'$ and $\partial \mathcal{J}/\partial \boldsymbol{p}^*$). Panel (B) shows the penalty contributions of the augmented Lagrangian as a function of constraint violation $c_i = 1 - \sum_k p^*_{ki}$. The left facet displays the linear multiplier term $\lambda c_i$, and the right facet shows the quadratic penalty $(\gamma/2)c_i^2$, each evaluated for multiple values of $\lambda$ and $\gamma$, respectively.

under both cumulative and isolated missing mass conditions, confirming that reference incompleteness constitutes a general challenge for all evaluated approaches, regardless of their underlying algorithmic design.

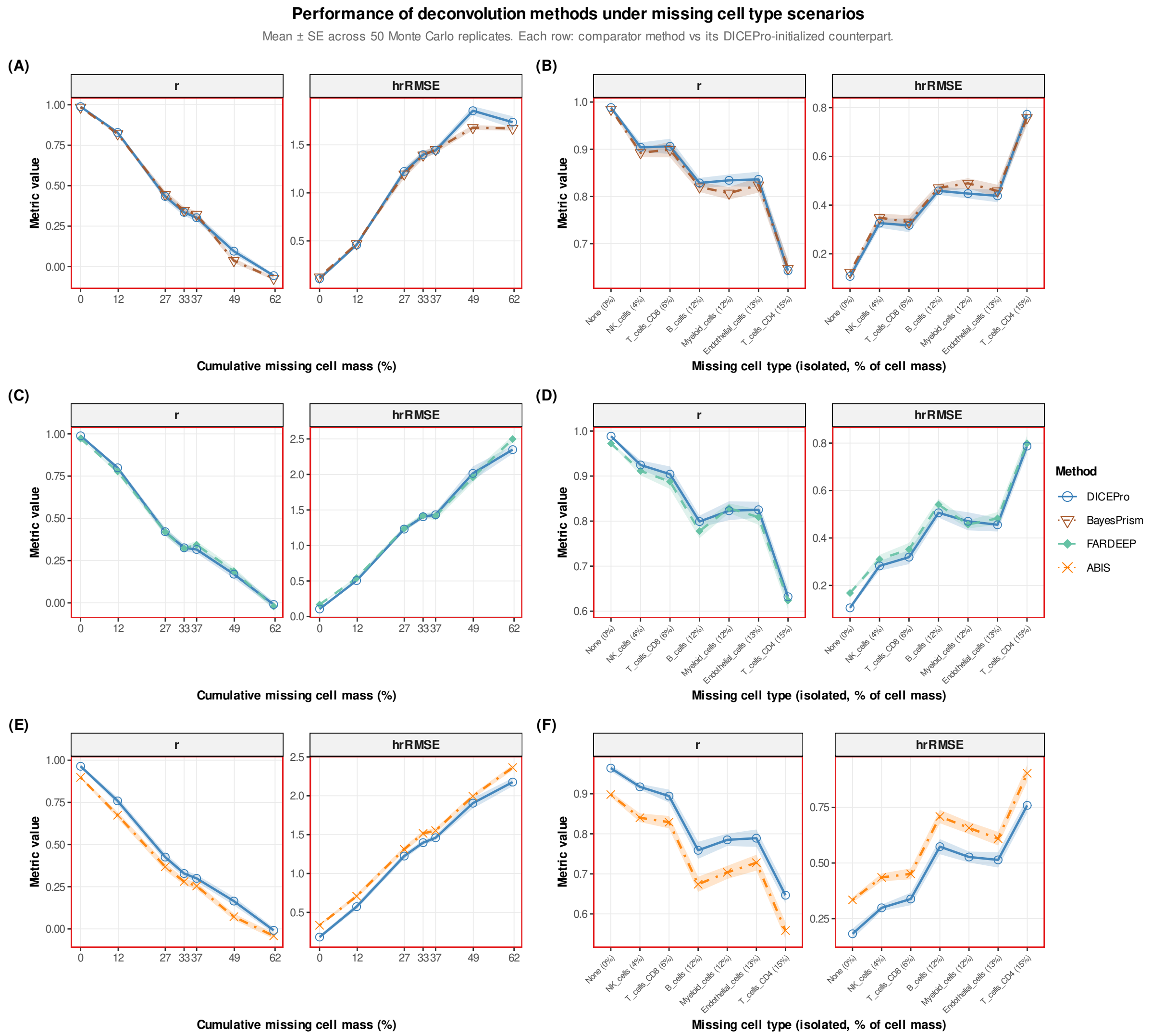

Figure S3: **Performance of BayesPrism, FARDEEP, and ABIS compared with the corresponding DICEPro-initialized methods under missing cell type scenarios.** Panels (A), (C), and (E) show cumulative missing mass conditions, where cell type groups are progressively removed from the reference matrix in the following order: B cells, CD4$^+$ T cells, CD8$^+$ T cells, NK cells, myeloid cells, and endothelial cells; panels (B), (D), and (F) show isolated missing mass conditions, where each cell type group is excluded individually from the reference matrix. In both scenarios, deconvolution was performed using the reduced reference matrix and predictions were compared to true proportions across 50 Monte Carlo replicates. Performance is evaluated by Pearson correlation coefficient ($r$) and hierarchical Relative Root Mean Square Error ($hrRMSE$). Ribbons indicate mean ± standard error across 50 Monte Carlo replicates. Each row contrasts a comparator method (BayesPrism, FARDEEP, or ABIS) with its DICEPro-initialized counterpart, sharing the same reduced reference signature matrix.

Against this background, DICEPro-initialized variants remained competitive across the full range of missing mass scenarios, generally matching or exceeding the performance of their corresponding base methods. The relative benefit of DICEPro initialization was particularly apparent under more severe missing mass conditions, where the post-deconvolution optimization framework appeared to partially compensate for the degraded reference information by imposing additional structure on the estimated proportion profiles. Under isolated missing mass conditions, where the impact of a single missing cell type group could be assessed independently, the DICEPro-initialized variants consistently maintained higher or equivalent $r$ values compared to their base method counterparts in the majority of excluded cell type groups, with no systematic deterioration attributable to the initialization step.

More strikingly, direct comparison of DICEPro variants initialized with BayesPrism, FARDEEP, and ABIS revealed a high degree of consistency between initialization strategies (Supplementary Figure S4). Despite the substantial methodological differences among these algorithms — ranging from Bayesian inference to constrained least squares and reference-based regression — all DICEPro variants produced remarkably similar performance profiles under both cumulative and isolated conditions. Performance differences between initialization strategies were negligible in relation to the general magnitude of degradation induced by missing cell types, indicating that the choice of initialization algorithm had little influence on the final result of deconvolution. Even when initialized from a comparatively weaker base method, `DICEPro` performance remained highly stable and consistently comparable to the best-performing approaches across all evaluated scenarios. Taken together, these observations suggest that the robustness of `DICEPro` is an intrinsic property of its optimization framework rather than a consequence of any particular initialization strategy, and that the post-deconvolution correction it applies is largely invariant to the starting point of the optimization.

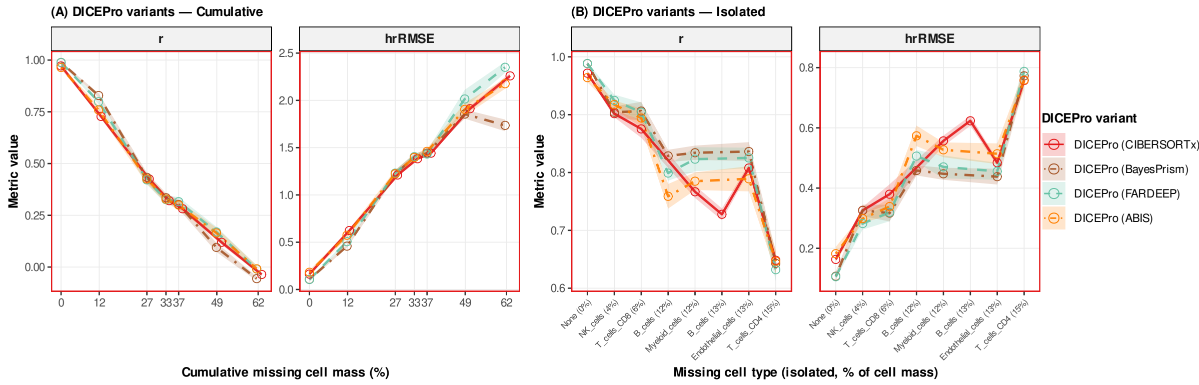

Figure S4: **Comparison of DICEPro performance across different initialization strategies (BayesPrism, FARDEEP, and ABIS) under missing cell type scenarios.** Panel (A) shows cumulative missing mass conditions, where cell type groups are progressively removed from the reference matrix in the following order: B cells, CD4$^+$ T cells, CD8$^+$ T cells, NK cells, myeloid cells, and endothelial cells; panel (B) shows isolated missing mass conditions, where each cell type group is excluded individually from the reference matrix. In both scenarios, deconvolution was performed using the reduced reference matrix and predictions were compared to true proportions across 50 Monte Carlo replicates. Performance is evaluated by Pearson correlation coefficient ($r$) and hierarchical Relative Root Mean Square Error ($hrRMSE$).

#### S1.5 Multi-objective optimization and Pareto front construction in DICEPro

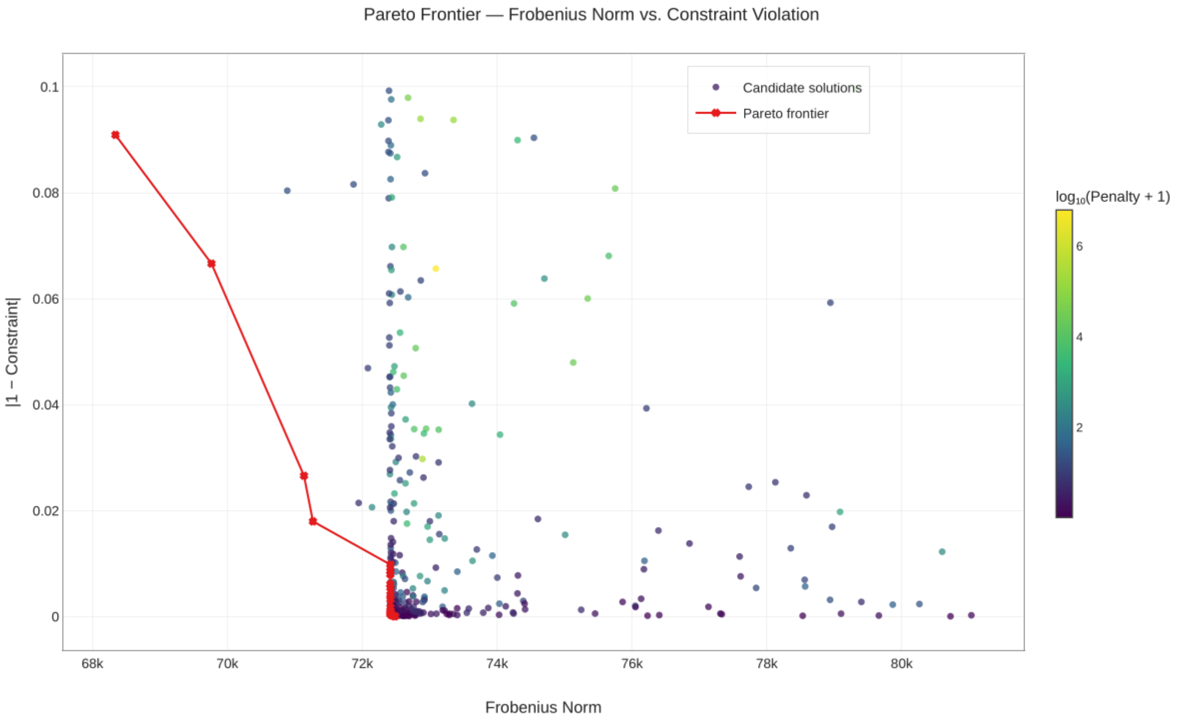

Figure S5: **Pareto frontier of DICEPro optimization: trade-off between reconstruction fidelity and constraint satisfaction.** Each point represents a candidate solution obtained during the hyperparameter search, characterized by its Frobenius norm (x-axis, measuring reconstruction error) and its deviation from the unit-sum constraint $|1 - \sum_k \hat{p}_k|$ (y-axis). The color encodes the $\log_{10}$-transformed penalty value, reflecting the degree of constraint violation incurred during optimization. The Pareto frontier (red line) identifies the set of non-dominated solutions that simultaneously minimize reconstruction error and constraint violation, with no candidate solution able to improve on both objectives at once. The concentration of Pareto-optimal solutions at low constraint deviation values demonstrates that `DICEPro` effectively enforces biological plausibility of proportion estimates while maintaining competitive reconstruction fidelity.

A critical but often overlooked limitation of standard deconvolution approaches is that minimizing reconstruction error alone — typically measured by the Frobenius norm between the observed bulk expression matrix and its estimated decomposition — is insufficient to guarantee biologically meaningful solutions. Indeed, unconstrained minimization of reconstruction error can yield cell-type proportion estimates that fall outside the biological simplex, producing negative values or proportions whose sum deviates substantially from unity. Although such solutions may achieve low reconstruction error, they remain biologically implausible, as cell-type proportions must be non-negative and sum to one within each sample.

`DICEPro` addresses this issue by reformulating deconvolution as a multi-objective optimization problem in which reconstruction fidelity and satisfaction of the unit-sum constraint are treated as distinct, potentially competing objectives. Rather than combining these objectives into a single scalar loss through fixed weighting — which would require arbitrary specification of their relative importance — `DICEPro` explicitly characterizes the trade-off between them using a Pareto front formulation. For each candidate solution in the hyperparameter space, optimization yields a point in a two-dimensional objective space composed of the Frobenius norm and the constraint deviation $|1 - \sum_k \hat{p}_k|$. The Pareto frontier is then defined as the set of non-dominated solutions, i.e., solutions for which no alternative simultaneously achieves lower reconstruction error and lower constraint violation (Figure S5).

Inspection of the Pareto frontier reveals several important properties of the `DICEPro` optimization landscape. First, the frontier is concentrated near low constraint deviation values, indicating that Pareto-optimal solutions satisfy the unit-sum constraint with high accuracy with most points achieving $|1 - \sum_k \hat{p}_k| \approx 0$. This suggests that enforcing biological plausibility does not necessarily compromise reconstruction fidelity and that a substantial region of the hyperparameter space yields solutions that are both accurate and biologically consistent. Second, solutions associated with high penalty values — reflecting strong constraint violations during optimization, as indicated by the color scale in Supplementary Figure S5 — are excluded from the Pareto frontier, demonstrating that the multi-objective formulation naturally disfavors biologically implausible solutions. Third, the frontier exhibits a characteristic elbow shape, identifying a region of diminishing returns in which further reductions in reconstruction error are achieved only at the expense of increased constraint violation. The selected solution corresponds to the knee point of this frontier, representing the most balanced trade-off between the two objectives.

Taken together, these results demonstrate that the multi-objective formulation implemented in `DICEPro` provides a principled and interpretable framework for balancing reconstruction fidelity and biological constraint satisfaction. More broadly, the Pareto front construction offers an effective strategy for automated hyperparameter selection without requiring arbitrary weighting between competing objectives.

### S2 Benchmarked deconvolution methods overview

This section describes the supervised deconvolution methods included in the benchmark. The methods span four methodological families: i) **constrained regression and quadratic programming** (`ABIS` (Monaco et al., 2019), `AutoGeneS` (Aliee and Theis, 2021), `DeconRNASeq` (Gong and Szustakowski, 2013), `EPIC` (Racle et al., 2017), `LinDeconSeq` (Li et al., 2020), `QProg` and `QProgwc` (Gong et al., 2011)); ii) **robust and regularized regression** (`CIBERSORTx` (Newman et al., 2019), `DCQ` (Altboum et al., 2014), `FARDEEP` (Hao et al., 2019), `RLR` and `RLS` (Sturm et al., 2019)); iii) **probabilistic and Bayesian modeling** (`BayesPrism` (Chu et al., 2022)); and iv) **machine learning-based frameworks** (`DISSECT` (Khatri et al., 2024)). All methods were applied under identical experimental conditions and their implementation details are provided in the subsections below.

#### S2.1 Constrained regression and optimization-based approaches

**`ABIS` (Absolute Immune Signal):** Monaco et al. (2019) introduce an explicit normalization for transcriptional output per cell type. Reference signatures are scaled by the estimated total mRNA content of each cell type before regression, so that the resulting estimates reflect actual cell abundances rather than relative transcript fractions. This normalization is particularly important for immune populations with large differences in transcriptional activity (e.g., plasma cells versus naive lymphocytes).

`AutoGeneS`: Aliee and Theis (2021) propose a data-driven gene selection framework that decouples feature selection from proportion estimation and requires no prior knowledge of marker genes. Rather than relying on literature-curated signatures or single-criterion tests (e.g., pairwise *t*-tests), `AutoGeneS` formulates gene selection as a multi-objective optimization problem: it simultaneously minimizes inter-cell-type correlation (measured by cosine similarity) and maximizes inter-cell-type distance (measured by Euclidean distance) over the space of candidate gene subsets. Because these two objectives are generally conflicting, no single gene set optimizes both simultaneously; the algorithm therefore searches for the Pareto-optimal frontier — the set of solutions not dominated by any other explored solution — using an evolutionary multi-objective optimizer. A final gene set is selected from this Pareto front by retaining the solution with the lowest mean pairwise correlation coefficient across cell types. The resulting signature matrix is then passed to a built-in regression module (NNLS by default) to estimate cell-type proportions. `AutoGeneS` accepts reference profiles from both sorted bulk RNA-seq and scRNA-seq data, making it compatible with modern single-cell atlases where canonical marker lists may not yet be established for novel subtypes.

`DeconRNASeq`: Gong and Szustakowski (2013) model bulk expression as a linear combination of reference profiles and solve a constrained non-negative least squares (NNLS) problem by quadratic programming. The method minimizes the residual sum of squares between observed bulk profiles and their estimated decompositions under the constraint that all fractions remain non-negative. Gene expression data must be supplied in linear scale, as the underlying mixture model is linear and log-transformation would violate this assumption. Proportions are renormalized to sum to one after estimation. The method requires a pre-specified reference signature matrix and is designed primarily for RNA-seq data, though it has been applied to microarray profiles after appropriate normalization.

`EPIC`: Racle et al. (2017) extend constrained least-squares regression with two biologically motivated adjustments. First, it incorporates cell-type-specific mRNA scaling factors that correct for differences in total RNA yield across cell populations, analogous to the approach of `ABIS` but estimated from independent reference datasets. Second, it explicitly reserves a fraction of the mixture for uncharacterized or unlisted cell types by introducing a catch-all unknown component, whose proportion is inferred alongside the reference cell types. This design makes `EPIC` particularly well suited to complex tissues such as the tumor microenvironment, where non-immune stromal and malignant populations are often absent from standard immune reference matrices.

`LinDeconSeq`: Li et al. (2020) decouple the selection of marker genes from proportion estimation. In the first step, each gene receives a cell-type specificity score quantifying how exclusively it is expressed in one cell type relative to all others; genes exceeding a specificity threshold are retained as markers. In the second step, weighted robust linear regression (RLM) is applied to the selected gene subset, where the weights are derived from the inverse of within-cell-type expression variance, reducing the influence of high-variance or unreliable genes on the final estimates.

`QProg` & `QProgwc`: Gong et al. (2011) proposed two quadratic programming deconvolution methods derived from a common constrained optimization framework. `QProg` solves the standard non-negative least squares problem without an explicit simplex constraint, whereas `QProgwc` extends this formulation by additionally enforcing a sum-to-one constraint on the estimated proportions alongside non-negativity, yielding fractions that form a valid probability simplex. The sum-to-one constraint in `QProgwc` makes it more appropriate for settings where the reference matrix is assumed to span all major cell populations present in the mixture.

#### S2.2 Robust and regularized regression approaches

**`CIBERSORTx`:** Newman et al. (2019) extend the original CIBERSORT framework by applying linear-kernel $\nu$-support vector regression ($\nu$-SVR) to estimate cell-type proportions. $\nu$-SVR concentrates optimization on the most informative genes (support vectors) and is inherently insensitive to genes that lie within the $\varepsilon$-tube around the regression hyperplane, providing natural robustness to irrelevant features. The parameter $\nu \in \{0.25, 0.50, 0.75\}$ controls the upper bound on the fraction of support vectors and the lower bound on the fraction of margin errors; the optimal value is selected as the one minimizing the resulting $\varepsilon$ (i.e., the width of the insensitive tube around the fitted hyperplane). `CIBERSORTx` adds two key capabilities: (i) automated construction of a signature matrix directly from scRNA-seq profiles, bypassing the need for pre-sorted bulk references, and (ii) cross-platform batch correction in two modes — S-mode, which adjusts the signature matrix to account for technical variation introduced by UMI-based or droplet-based single-cell platforms (e.g., 10x Chromium), and B-mode, which adjusts the bulk mixture data to correct for cross-platform differences with the reference. It can optionally impute cell-type-specific expression profiles at the group level (averaged across all samples) or at high resolution (per sample), enabling downstream analyses of context-specific transcriptional programs.

**`DCQ` (Digital Cell Quantifier):** Altboum et al. (2014) apply elastic-net regularized linear regression, combining $\ell_1$ (lasso) and $\ell_2$ (ridge) penalties to simultaneously perform feature selection and coefficient shrinkage. The $\ell_1$ component drives the coefficients of irrelevant marker genes to zero, while the $\ell_2$ component handles correlated markers by distributing weight across them rather than arbitrarily selecting one. `DCQ` uses a set of curated immune marker genes and stabilizes estimates through an ensemble strategy: multiple regression instances are fit to resampled gene subsets and predictions are averaged across the ensemble to reduce sensitivity to individual noisy measurements.

**`FARDEEP` (Fast And Robust DEconvolution of Expression Profiles):** Hao et al. (2019) addresses the distortion of proportion estimates caused by outlier genes through an adaptive least-trimmed squares (aLTS) algorithm. Unlike standard LTS, which discards a fixed proportion of observations, aLTS iteratively identifies and removes observations whose residuals exceed a data-driven threshold, computed from the empirical residual distribution at each iteration. This adaptive trimming ensures convergence in finite steps regardless of the true outlier fraction, and the algorithm guarantees a globally optimal solution under its trimming criterion. After outlier removal, a standard NNLS problem is solved on the retained genes. `FARDEEP` produces both relative proportions and an estimate of a scaling factor related to absolute cell abundance, and has been validated on microarray (LM22) and RNA-seq references.

**`RLR` (Robust Linear Regression) & `RLS` (Robust Least Squares):** Sturm et al. (2019) introduce two robustness-adjusted wrappers around standard linear regression. `RLR` applies Huber M-estimators in an iteratively reweighted least squares (IRLS) procedure: at each iteration, observations (genes) are assigned weights inversely proportional to their absolute residual when the residual exceeds a threshold $k$ (typically $k = 1.345\hat{\sigma}$ for 95% asymptotic efficiency under normality), while observations with small residuals retain full weight. This down-weighting progressively reduces the influence of genes with aberrant expression without completely discarding them, preserving power when the fraction of outliers is moderate. `RLS` instead augments the least-squares objective with leverage-based corrections, reducing the influence of high-leverage observations that can disproportionately determine the regression hyperplane. Both methods were evaluated as part of the benchmarking framework of Sturm et al. (2019).

#### S2.3 Probabilistic and Bayesian modeling approaches

**`BayesPrism`:** Chu et al. (2022) propose a fully Bayesian deconvolution framework that models gene expression within each cell type using a Dirichlet-multinomial distribution, with scRNA-seq profiles providing informative priors on cell-type-specific expression. The joint posterior distribution over cell-type fractions $\theta$ and cell-type-specific expression profiles $Z$ is:

$$P(\theta, Z \mid X, \phi) \propto P(X \mid \theta, Z)\, P(Z \mid \phi)\, P(\theta),$$

where $X$ denotes the observed bulk counts, $\phi$ the reference scRNA-seq profiles, and $P(\theta)$ a symmetric Dirichlet prior. To accommodate within-cell-type transcriptional heterogeneity (e.g., M1/M2 macrophage polarization or cell-cycle variation in malignant cells), `BayesPrism` introduces a cell-state layer: scRNA-seq clusters are treated as fine-grained states nested within cell types, and the posterior is marginalized over states to recover cell-type-level proportions. Posterior inference is performed via Gibbs sampling. An optional second step updates the reference $\phi$ using a posteriori estimates — either per-sample maximum likelihood or cross-sample maximum a posteriori estimation — improving robustness when reference-mixture domain shifts are expected.

#### S2.4 Machine learning-based approaches

**`DISSECT`:** Khatri et al. (2024) propose a semi-supervised deep learning framework specifically designed to overcome the domain shift between synthetically generated training data and real bulk RNA-seq measurements — a fundamental limitation shared by purely supervised approaches such as `Scaden` (Menden et al., 2020). Its architecture relies on two components: (i) an ensemble of multilayer perceptrons (MLPs) with architecture Input $\rightarrow$ ReLU6(512) $\rightarrow$ ReLU6(256) $\rightarrow$ ReLU6(128) $\rightarrow$ ReLU6(64) $\rightarrow$ Softmax($K$) for estimating the proportions of the $K$ cell types, and (ii) a conditional autoencoder for estimating cell-type-specific expression profiles, both contributing to the training objective. Training minimizes a combined loss:

$$\mathcal{L} = \mathcal{L}_{\text{supervised}}(p_{\text{sim}}) + \alpha\, \mathcal{L}_{\text{consistency}}(p_{\text{real}}),$$

where $\mathcal{L}_{\text{supervised}}$ is the proportional prediction loss on simulated pseudo-bulk data and $\mathcal{L}_{\text{consistency}}$ penalizes prediction instability in real bulk samples under data augmentation, acting as an unsupervised domain adaptation signal. The balance parameter $\alpha$ controls the strength of domain adaptation. Consistency regularization encourages the model to produce stable and biologically plausible predictions when applied to real bulk data that may deviate from the training distribution.

### S3 Hierarchical performance metrics for deconvolution evaluation

To evaluate deconvolution performance, we model true and estimated cell-type proportions using a hierarchical linear framework that decomposes variability into population-level, sample-level, and residual components. This multilevel formulation allows variance to be partitioned into biologically significant sources, providing a basic basis for performance assessment that avoids biases introduced by treating proportion estimates as independent observations (Gelman and Hill, 2006; McCulloch et al., 2008).

#### Hierarchical model

Let $p_{ij}$ denote the true proportion of the cell population $j \in \{1, \ldots, K\}$ in sample $i \in \{1, \ldots, N\}$, and let $\hat{p}_{ij}$ be the corresponding estimated proportion.

##### Standard formulations

The intraclass correlation coefficient `ICC(3,1)`, as defined by Shrout and Fleiss (1979), assumes a two-way mixed-effects model and decomposes the total variance into three components

$$\texttt{ICC(3,1)} = \frac{\sigma_r^2}{\sigma_r^2 + \sigma_c^2 + \sigma_e^2},$$

where $\sigma_r^2$ is the variance attributable to subjects (variability between-rows), $\sigma_c^2$ captures systematic measurement bias (variability between-columns), and $\sigma_e^2$ is the residual error. Reliability is conventionally interpreted as poor ($< 0.50$), moderate (0.50–0.75), good (0.75–0.90) or excellent ($> 0.90$) (Koo and Li, 2016). The standard root mean square error between the observed and predicted proportions is

$$\texttt{rmse} = \sqrt{\frac{1}{NK} \sum_{i=1}^{N} \sum_{j=1}^{K} \left(p_{ij} - \hat{p}_{ij}\right)^2}.$$

##### Adapted formulations

Because cell-type proportions exhibit a nested structure — samples are measured across multiple cell populations — a simple two-group decomposition is insufficient. Therefore, both observed and predicted proportions are modeled under the same hierarchical linear structure (Bates et al., 2015):

$$y_{ijk} = \mu + u_i + v_{ij} + \epsilon_{ijk},$$

where $y_{ijk}$ represents either $p_{ijk}$ or $\hat{p}_{ijk}$; $\mu$ is the global mean proportion; $u_i \sim \mathcal{N}(0, \sigma^2_{\text{population}})$ captures the variability between cell populations; $v_{ij} \sim \mathcal{N}(0, \sigma^2_{\text{sample}})$ represents sample-specific deviations nested within each cell population; and $\epsilon_{ijk} \sim \mathcal{N}(0, \sigma^2_e)$ denotes residual prediction error. The total biological variance is

$$V_T = \sigma^2_{\text{population}} + \sigma^2_{\text{sample}}.$$

The model parameters are estimated using restricted maximum likelihood (REML), with variance components extracted from the fitted model (Bates et al., 2015).

Three complementary metrics are derived from this hierarchical variance decomposition, each attaining its optimal value under perfect prediction ($\sigma^2_e = 0$).

The adapted `ICC3` generalizes `ICC(3,1)` by replacing the simple $(\sigma^2_r, \sigma^2_c)$ decomposition with the nested biological variance $V_T$:

$$\texttt{ICC3} = \frac{V_T}{V_T + \sigma^2_e} = \frac{\sigma^2_{\text{population}} + \sigma^2_{\text{sample}}}{\sigma^2_{\text{population}} + \sigma^2_{\text{sample}} + \sigma^2_e} \in [0, 1],$$

with $\texttt{ICC3} = 1$ indicating perfect agreement and $\texttt{ICC3} = 0$ indicating that all variance is residual.

The concordance correlation coefficient (LI, 1989) adapted to the hierarchical setting combines the same variance components:

$$\texttt{CCC} = \frac{2\,V_T}{2\,V_T + \sigma^2_e} \in [0, 1],$$

with $\texttt{CCC} = 1$ under perfect prediction. This formulation assumes that the systematic bias between observed and predicted proportions is absorbed by the fixed effects of the mixed model, leaving $\sigma^2_e$ as the sole disagreement term.

The hierarchical relative root mean square error (`hrRMSE`) is the scale-free counterpart to `rmse`, anchored to the biological signal rather than raw proportion values:

$$\texttt{hrRMSE} = \sqrt{\frac{\sigma^2_e}{V_T}} \in [0, +\infty),$$

with $\texttt{hrRMSE} = 0$ indicating perfect prediction. Although unbounded above, values below 1 indicate that the residual prediction error is smaller than the biological signal ($\sigma^2_e < V_T$), providing a natural interpretability threshold.

The three adapted metrics are mutually consistent and satisfy the following closed-form relationships:

$$\texttt{ICC3} = \frac{V_T}{V_T + \sigma^2_e}, \qquad \texttt{CCC} = \frac{2\,V_T}{2\,V_T + \sigma^2_e}, \qquad \texttt{hrRMSE} = \sqrt{\frac{\sigma^2_e}{V_T}},$$

$$\texttt{CCC} = \frac{2\,\texttt{ICC3}}{1 + \texttt{ICC3}}, \qquad \texttt{hrRMSE}^2 = \frac{1 - \texttt{ICC3}}{\texttt{ICC3}}.$$

As $\sigma^2_e \to 0$: $\texttt{ICC3} \to 1$, $\texttt{CCC} \to 1$, $\texttt{hrRMSE} \to 0$.

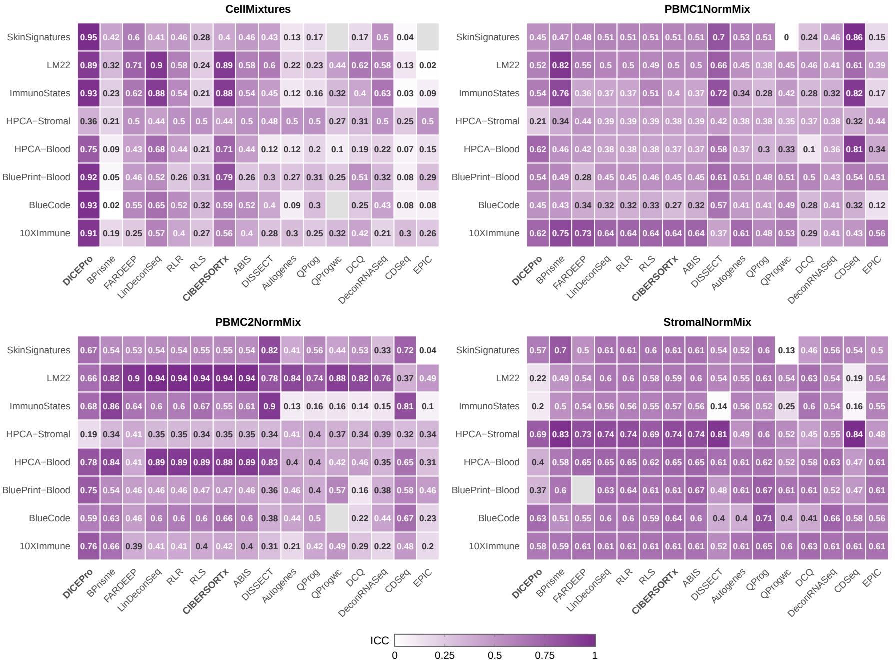

**(a)** Intraclass Correlation Coefficient (ICC3)

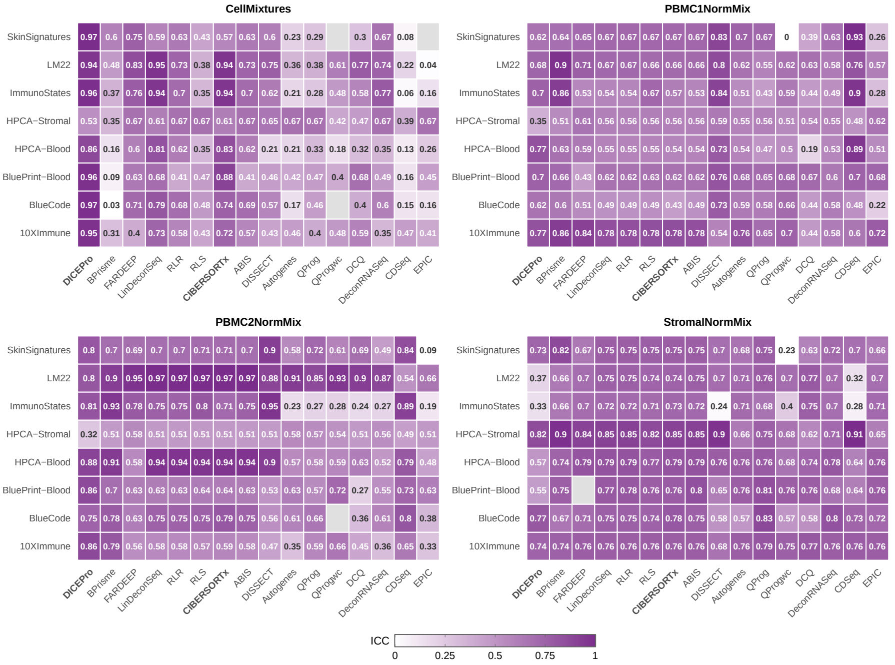

**(b)** Concordance Correlation Coefficient (CCC)

Figure S6: **Hierarchical agreement metrics on real bulk RNA-seq datasets.** Heatmaps of (a) Intraclass Correlation Coefficient (ICC3) and (b) Concordance Correlation Coefficient (CCC) for all benchmark methods across four datasets (CellMixtures, PBMC1NormMix, PBMC2NormMix and StromalNormMix) and eight reference matrices. Both metrics are derived from the hierarchical variance decomposition and range in $[0, 1]$. Darker shading indicates better performance in both panels: higher ICC3 and higher CCC.

Andrew Gelman and Jennifer Hill. *Data Analysis Using Regression and Multilevel/Hierarchical Models*. Analytical Methods for Social Research. Cambridge University Press, 2006.

T Gong and JD Szustakowski. Deconrnaseq: a statistical framework for deconvolution of heterogeneous tissue samples based on mrna-seq data. *Bioinformatics*, 29:1083–1085, 2013. doi: 10.1093/bioinformatics/btt090.

T Gong, N Hartmann, IS Kohane, V Brinkmann, F Staedtler, M Letzkus, S Bongiovanni, and JD Szustakowski. Optimal deconvolution of transcriptional profiling data using quadratic programming with application to complex clinical blood samples. *PLoS One*, 6(11):e27156, 2011. doi: 10.1371/journal.pone.0027156.

Y Hao, M Yan, BR Heath, YL Lei, and Y Xie. Fast and robust deconvolution of tumor infiltrating lymphocyte from expression profiles using least trimmed squares. *PLoS Comput Biol*, 15(5):e1006976, 2019. doi: 10.1371/journal.pcbi.1006976.

Robin Khatri, Pierre Machart, and Stefan Bonn. Dissect: deep semi-supervised consistency regularization for accurate cell type fraction and gene expression estimation. *Genome Biology*, 25(1):112, 2024. doi: 10.1186/s13059-024-03251-5.

Diederik P Kingma and Jimmy Ba. Adam: A method for stochastic optimization. In *Proceedings of the 3rd International Conference on Learning Representations (ICLR)*, San Diego, CA, 2015. arXiv:1412.6980.

Terry K. Koo and Mae Y. Li. A guideline of selecting and reporting intraclass correlation coefficients for reliability research. *Journal of Chiropractic Medicine*, 15(2):155–163, 2016. doi: 10.1016/j.jcm.2016.02.012.

Huamei Li, Amit Sharma, Wenglong Ming, Xiao Sun, and Hongde Liu. A deconvolution method and its application in analyzing the cellular fractions in acute myeloid leukemia samples. *BMC Genomics*, 21:652, 2020. doi: 10.1186/s12864-020-06888-1.

Lin LI. A concordance correlation coefficient to evaluate reproducibility. *Biometrics*, 45 (1):255–68, Mar 1989.

Dong C Liu and Jorge Nocedal. On the limited memory BFGS method for large scale optimization. *Math Program*, 45(1–3):503–528, 1989. doi: 10.1007/BF01589116.

Charles E. McCulloch, S. R. Searle, and John M. Neuhaus. *Generalized, linear, and mixed models / Charles E. McCulloch, Shayle R. Searle, John M. Neuhaus.* Wiley series in probability and statistics. Wiley, Hoboken, N.J, 2nd ed. edition, 2008. ISBN 9780470073711.

Kevin Menden, Matteo Marouf, Sergio Oller, Anupriya Dalmia, Daniel Sumner Magruder, Karin Kloiber, Peter Heutink, and Stefan Bonn. Deep learning-based cell composition analysis from tissue expression profiles. *Science Advances*, 6(30):eaba2619, 2020. doi: 10.1126/sciadv.aba2619.

G Monaco, B Lee, W Xu, S Mustafah, YY Hwang, C Carre, N Burdin, L Visan, M Ceccarelli, M Poidinger, A Zippelius, JP de Magalhaes, and A Larbi. Rna-seq signatures normalized by mrna abundance allow absolute deconvolution of human immune cell types. *Cell Rep*, 26(6):1627–1640.e7, 2019. doi: 10.1016/j.celrep.2019.01.041.

Brian B Nadel, Meritxell Oliva, Benjamin L Shou, Keith Mitchell, Feiyang Ma, Dennis J Montoya, Alice Mouton, Sarah Kim-Hellmuth, Barbara E Stranger, Matteo Pellegrini, and Serghei Mangul. Systematic evaluation of transcriptomics-based deconvolution methods and references using thousands of clinical samples. *Briefings in Bioinformatics*, 22(6):bbab265, 11 2021. doi: 10.1093/bib/bbab265.

Aaron M Newman, Chih Long Liu, Michael R Green, Andrew J Gentles, Weiguo Feng, Yue Xu, Chuong D Hoang, Maximilian Diehn, and Ash A Alizadeh. Robust enumeration of cell subsets from tissue expression profiles. *Nature methods*, 12(5):453–457, 2015.

AM Newman, CB Steen, CL Liu, AJ Gentles, AA Chaudhuri, F Scherer, MS Khodadoust, MS Esfahani, BA Luca, D Steiner, M Diehn, and AA Alizadeh. Determining cell type abundance and expression from bulk tissues with digital cytometry. *Nat Biotechnol*, 37 (7):773–782, 2019. doi: 10.1038/s41587-019-0114-2.

Jorge Nocedal. Updating quasi-Newton matrices with limited storage. *Math Comput*, 35 (151):773–782, 1980. doi: 10.1090/S0025-5718-1980-0572855-7.

J Racle, K de Jonge, P Baumgaertner, DE Speiser, and D Gfeller. Simultaneous enumeration of cancer and immune cell types from bulk tumor gene expression data. *eLife*, 6: e26476, 2017. doi: 10.7554/eLife.26476.

PE Shrout and JL Fleiss. Intraclass correlations: uses in assessing rater reliability. *Psychol Bull*, 86(2):420–428, 1979. doi: 10.1037//0033-2909.86.2.420.

G Sturm, F Finotello, F Petitprez, JD Zhang, J Baumbach, WH Fridman, M List, and T Aneichyk. Comprehensive evaluation of transcriptome-based cell-type quantification methods for immuno-oncology. *Bioinformatics*, 35(14):i436–i445, 2019. doi: 10.1093/bioinformatics/btz363.
